# Arf GTPase-activating protein ASAP1 specifically binds to 440-kD ankyrin-B

**DOI:** 10.1101/2024.01.05.574302

**Authors:** Yubing Li, Yipeng Zhao, Yaojun He, Mingjie Zhang, Keyu Chen

**Affiliations:** Greater Bay Biomedical Innocenter, Shenzhen Bay Laboratory, Shenzhen 518036, China; Division of Life Science, Hong Kong University of Science and Technology, Clear Water Bay, Kowloon, Hong Kong, China; School of Life Sciences, Southern University of Science and Technology, Shenzhen 518055, China

**Keywords:** ASAP1, ASAP2, ankyrin-B, ankyrin-G, SH3 domain, APC, MICAL1, Clasp1, Clasp2

## Abstract

The 440-kD giant ankyrin-B (gAnkB) exclusively localizes to axons and is essential for axon development. However, proteins that specifically bind to gAnkB but not to other isoforms of ankyrins are poorly understood. Here, we discovered that an Arf GTPase-activating protein ASAP1 and ASAP2 specifically binds to a short and disordered sequence unique to gAnkB. Biochemical studies showed that the SH3 domain of ASAP1 binds to a 12-residue, positively charged peptide fragment from gAnkB. The high-resolution structure of the ASAP1-SH3 domain in complex with the gAnkB peptide revealed the mechanism underlying this non-canonical SH3 domain-mediated target recognition. Further structural and bioinformatic analysis revealed additional previously unknown ASAP1-SH3 binding partners including Clasp1 and Clasp2, both of which are well-known microtubule regulators. Among all known ASAP1-SH3 binders including those identified in the current study, gAnkB has the strongest affinity in binding to ASAP1. Our results suggest that ASAP1 may function together with gAnkB in regulating axonal cytoskeletons.

## Introduction

Ankyrin-B is a member of the ankyrin scaffold protein family, which includes ankyrin-R, ankyrin-B, and ankyrin-G encoded by ANK1, ANK2, and ANK3 genes, respectively. These proteins have diverse but evolutionarily conserved functions in different tissues and cells^1^. Within neuronal axons, an extra-large isoform of ankyrin-B (440-kD) known as giant ankyrin-B (gAnkB) is expressed ^2^. This isoform is produced through alternative mRNA splicing, acquiring an extremely long exon encoding an intrinsically disordered insertion sequence called the neurospecific domain (Figure 1A) ^3^. Interestingly, ankyrin-G, encoded by another gene ANK3 and as a paralog of ankyrin-B, also expresses a giant splicing variant specifically targeted to axons. The two giant ankyrins display distinct sub-axonal localizations. Giant ankyrin-G (gAnkG) clusters at the axon initial segments (AIS), while gAnkB is excluded from AIS and localized throughout the rest of the unmyelinated axonal trunk in neurons ^4–6^. Both proteins share a list of membrane proteins and cytoskeletal structures as their binding partners using their highly homologous structured protein domains ^1,2,5,7,8^. However, the molecular mechanisms underlying the distinct axonal sub-compartment localizations and their specific functions in such sub-compartments for the two giant ankyrins are unknown. One possible reason is that the ∼2,000- 3,000-residue giant insertions in the two ankyrins are different and thus may contribute to the unique localizations and functions of gAnkB and gAnkG in axons. Furthermore, gAnkB is a high-risk factor for autism spectrum disorders (ASDs), and several de novo ASD mutations have been identified in the unstructured giant insertion region of the protein ^2,9,10^. gAnkB has been shown to repress stochastic axon branching through direct interaction with cortical axonal microtubules, and the loss of the interaction may contribute to the autism pathophysiology ^1,6,8^. However, whether and how the interaction between gAnkB and microtubules is dynamically regulated is not clear. In this study, we aimed to identify specific gAnkB binding proteins from mice brain by performing affinity purification combined with mass spectrometry using a fragment of gAnkB insertion as the bait. We identified an Arf GTPase-activating protein ASAP1 as a strong and specific binder of gAnkB.

**Figure 1.**
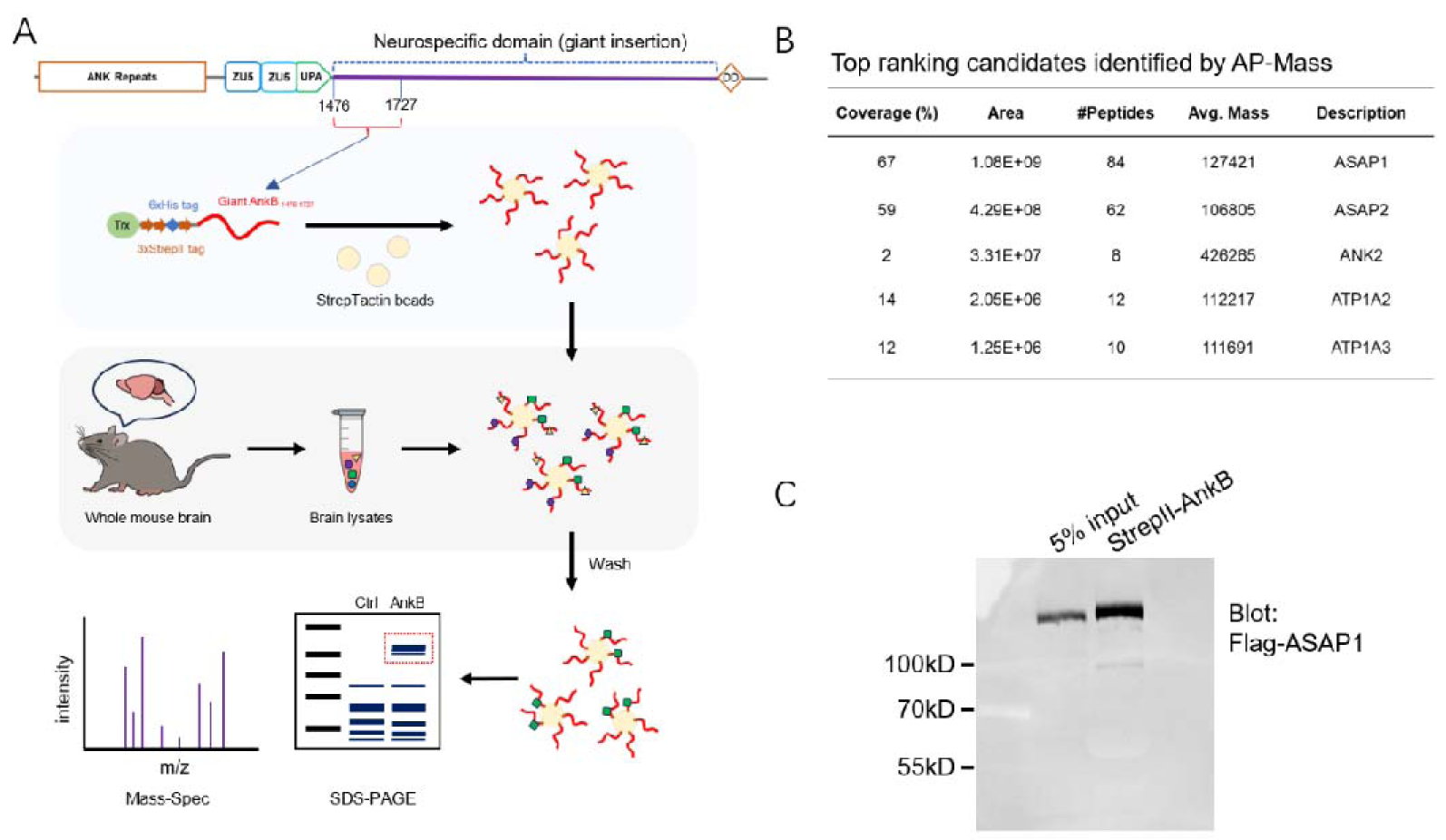
Affinity purification and mass spectrometry identification of gAnkB binding proteins. A: Illustration of the affinity purification process. A fragment (aa. 1476-1727) from gAnkB neurospecific domain was expressed in E. Coli and purified as a fusion protein with Trx-His_6_- 3×StrepII tag and used as a bait to pull down proteins from the mouse brain lysate. B: List of the top-ranked proteins identified by the affinity purification. “Area” means the highest peak intensity of the MS2 spectrum peak corresponding to the peptide segment; “Coverage%” means the percentage of protein sequences that can be covered by all the identified peptides. C: Pull-down assay of the binding between purified gAnkB (aa. 1476-1727) to Flag-tagged full-length ASAP1 expressed in HEK293T cells.

Arf GAP with SH3 Domain, Ankyrin Repeat and PH Domain 1 (ASAP1, also called AMAP1, DDEF1, DEF1, or Centaurin β4) is a PI_4,5_P_2_-dependent Arf GTPase-activating protein ^11–14^. ASAP1 can regulate membrane and cytoskeletal dynamics and is involved in controlling cell adhesive structures, including invadosomes ^11,13–16^, podosomes ^17–20^ and focal adhesions ^13,21,22^ during cell attachment and migration ^16,21,23–25^. The SH3 domain of ASAP1 is involved in the interactions with focal adhesion kinase (FAK), CD2AP, and Src ^12,14,18,26,27^; the N-terminal BAR domain functions in actin bundling and autoinhibition ^18,28^; the pleckstrin homology (PH) domain modulates the ARF-GAP activity ^28–30^. ASAP1 has also been implicated in promoting malignant phenotypes in cancers and defects in immune cells ^19,31–34^, its role in neuronal cells remains to be investigated.

SH3 domains are prevalent protein modules involved in various cellular processes such as intracellular signaling, cytoskeletal rearrangements, and immune responses ^35,36^. Human proteome contains around 300 SH3 domains ^37–40^. SH3 domains typically recognize proline-rich sequences containing a core “P-X-X-P” sequence element, where “X” represents any amino acid, through defined surface pockets ^41–44^. However, given that about 25% human proteins contain proline-rich sequences ^35^, the target binding specificities of SH3 domains must involve amino acid residues different from or outside of the “P-X-X-P” element . The interaction between ASAP1 SH3 and gAnkB described in the current study represents one such example.

## Results

### Identification of ASAP1 as a binding partner of gAnkB

In attempting to identify proteins that specifically bind to gAnkB but not to gAnkG, we used purified recombinant proteins corresponding to different fragments in the giant axon-specific disordered insertion as the baits for purifying binding proteins from mouse brain lysates. In this study, a 250-residue fragment of AnkB protein (aa. 1476-1727; Fig. 1A), tagged with three streptavidin-binding peptides (StrepII tags), was the bait proteins. Several distinct bands were detected from the SDS-PAGE of affinity purified proteins. Mass spectrometry analysis identified several top ranked candidates including ASAP1, ASAP2, ANK2, ATP1A2, and ATP1A3 (Fig. 1B). Among these, ANK2 (AnkB) should be a bait protein contaminate. ATP1A2/A3 are also likely non-specific binders due to their high abundance brain cells and their frequent recoveries when using other proteins as baits. ASAP1 and ASAP2 belong to the same family Arf GTPase-activating proteins and were recovered with very high peptide sequence coverages (Fig. 1B). Therefore, we tested the interaction between ASAP1 and gAnkB by performing pull-down experiments using purified 3×StrepII AnkB protein as the bait and flag-tagged ASAP1 full-length protein expressed in HEK293 cells. As shown in Figure 1C, the interaction between AnkB and the full-length ASAP1 was robustly detected.

### SH3 domain of ASAP1 binds to a short fragment of gAnkB

ASAP1 comprises a N-terminal BAR domain, a PH domain, an Arf-GAP domain, three ankyrin-repeats, followed by a C-terminal long-disordered region and an SH3 domain (Fig. 2A). To confirm the direct binding between ASAP1 and gAnkB and to determine the exact sites on both proteins responsible for the binding, we measured the binding affinity using isothermal titration calorimetry (ITC) with purified proteins. As depicted in Figure 2B, the SH3 domain (aa. 1083- 1147) of ASAP1 binds to gAnkB with a Kd ∼0.14 μM. Neither the proline-rich region (aa. 732- 915) nor the “E/DLPPKP”-repeat sequence (aa. 903-1082) displayed any interaction with AnkB. Therefore, the ASAP1 SH3 domain is responsible for binding to gAnkB. Similarly, to locate the ASAP1 SH3 binding sites on the gAnkB, we truncated the AnkB bait fragment (aa. 1477-1792) into shorter segments and determined their affinities with ASAP1 SH3 by ITC. As summarized in Figure 2B, the shortest sequence capable of binding ASAP1 was mapped to a 12-residue fragment of gAnkB (aa. 1699-1710). This short peptide is highly conserved in vertebrate gAnkB (Fig. S1A), but does not exist in gAnkB’s paralog ankyrin-G or ankyrin-R, suggesting that ASAP1 is a specific gAnkB binder in animal brains. We further note that the binding between ASAP1 and gAnkB is the strongest interaction among all reported ASAP1-SH3 binding proteins ^45^. In addition, strong binding between the SH3 domain of ASAP2, a homolog of ASAP1 that also identified in our AP-Mass, and gAnkB is also detected by ITC (Kd ∼68 nM; Fig. S1E).

**Figure 2.**
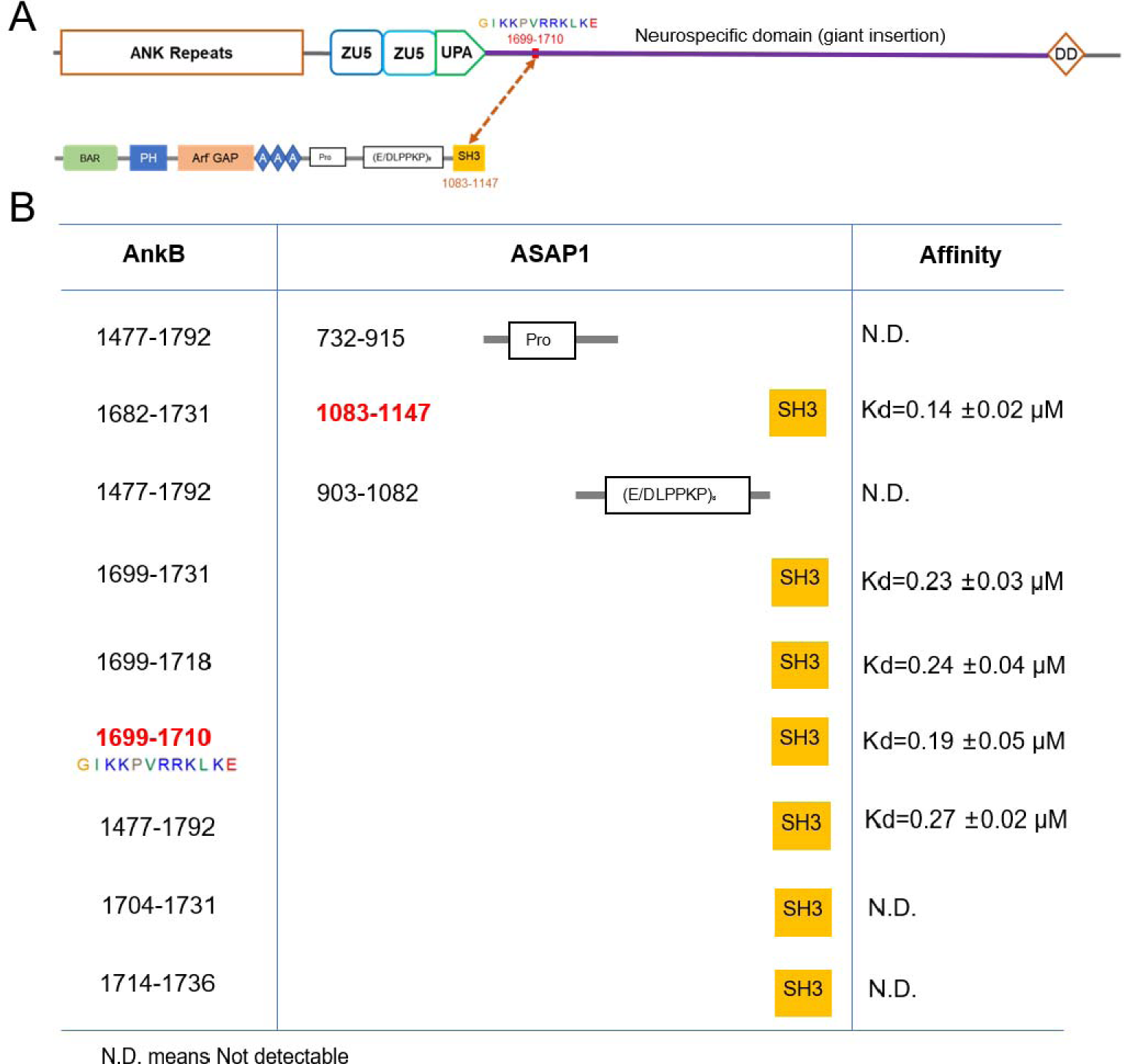
Mapping the interaction between gAnkB and ASAP1. A: Domain organizations of human 440-kD gAnkB and ASAP1. The minimal binding regions from the two proteins revealed from experiments in panel B are indicated by an arrow. B: Affinity directed binding site mapping of the interaction between gAnkB and ASAP1. Protein boundaries for each ITC-based binding reaction and derived dissociate coefficients are listed. Numbers labeled in red mean these are the shortest fragments in each protein that possess the full binding capability.

### Crystal structure of the ASAP1-SH3/AnkB complex reveals a new SH3 target binding mode

The binding of SH3 domains to their ligand peptides typically involves a minimal “P-X-X-P” motif. Curiously, the ASAP1 SH3 binding peptide from gAnkB contains only one proline residue, and yet the peptide binds to the SH3 domain with an affinity (Kd ∼0.14 μM) stronger than most of the reported SH3/target interactions. To elucidate the mechanism underlying the observed unusual interaction, we determined the crystal structure of ASAP1-SH3 in complex with the gAnkB peptide at the atomic resolution of 2.07 Å (Fig. 3A-B, Table 1).

**Figure 3.**
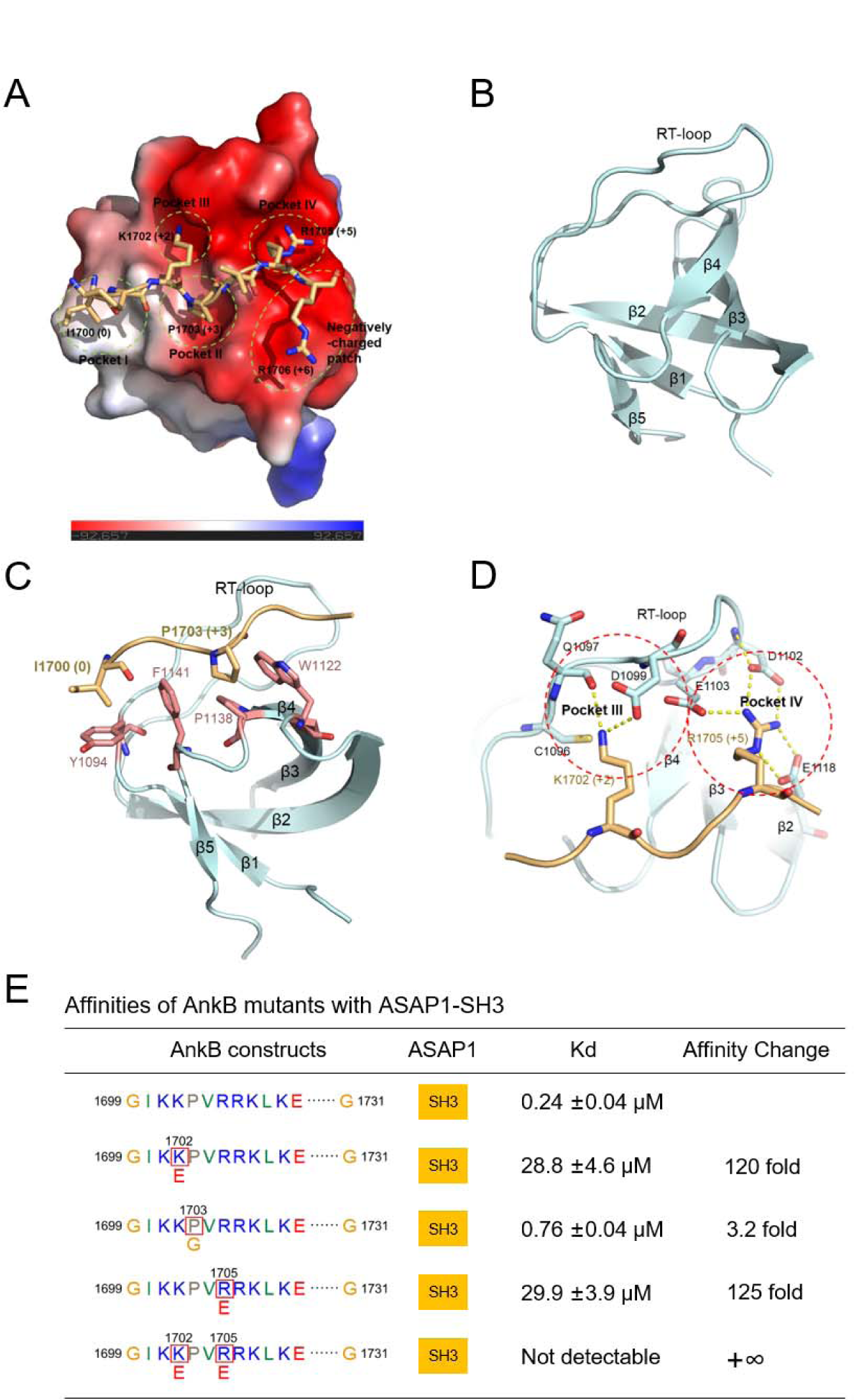
Four binding pockets and a negative-charged surface patch are identified from the gAnkB/ASAP1-SH3 crystal structure. A: Structure of ASAP1-SH3 in complex with the gAnkB peptide. ASAP1-SH3 is shown in surface charge potential model and the gAnkB peptide is shown in stick model. Four identified binding pockets on SH3 for AnkB binding are circled by green dished lines, labeled as Pocket I∼IV respectively. A highly negatively-charged surface patch on SH3 is also circled by green dished line. Numbers within the brackets indicates the designated position of amino acid residue on gAnkB (see Fig. 4D for detailed positions) B. Ribbon diagram of the ASAP1-SH3 domain in complex with an identical orientation as the surface model in panel A for locating key gAnkB binding elements from the SH3 domain (see Fig. 4C for detailed positions of amino acid residues in this domain). C: Ribbon diagram combined with the stick model showing the interact residues in pockets I & II in the complex. D: Stick model showing the charge-charge interactions between ASAP1-SH3 and the gAnkB peptide in pockets III & IV (see Fig. 4C & 4D for detailed positions of amino acid residues). E: Affinities of the gAnkB peptide mutants binding to ASAP1-SH3 determined by ITC.

**Table 1.** Statistics of X-ray Crystallographic Data Collection and Model refinement.

|  |  |
| --- | --- |
| Dara Sets |  |
| Space group | P 1 |
| Wavelength (Å) | 0.979 |
| Unit Cell parameters (Å) | a = 39.9225, b = 52.0739, c = 56.9089, $\alpha$ = 111.416, $\beta$ = 90.1631, $\gamma$ = 112.462 |
| Resolution range (Å) | 32.49 - 2.07 (2.12 - 2.07) |
| No. of unique reflections | 23173 (1695) |
| Redundancy | 3.2 (2.8) |
| I/$\sigma$ | 8.1 (5.7) |
| Completeness (%) | 97.6 (95.4) |
| R<sub>merge</sub>(%) | 12.1 (21.3) |
| Structure refinement |  |
| Resolution (Å) | 32.49 - 2.07 (2.63-2.07) |
| R<sub>cryst</sub>/R<sub>free</sub>(%) | 20.02 (22.14)/23.09 (28.36) |
| Rmsd bonds (Å)/angles (°) | 0.009/1.559 |
| Average B factor | 17.11 |
| No. of atoms |  |
| Protein atoms | 2226 |
| Water | 305 |
| Other molecules | 1 |
| No. of reflections |  |
| Working set | 23129 |
| Test set | 1132 |
| Ramachandran plot regions |  |
| Favored (%) | 98.06 |
| Allowed (%) | 1.55 |
| Outliers (%) | 0.39 |

In the complex structure, ASAP1-SH3 adopts the typical five-stranded SH3 fold. The ASAP1-SH3 also contains two SH3 domain-defining pockets (pockets I & II) responsible for binding two proline resides in the canonical “P-X-X-P” motif (Fig. 3A, 3C) ^43^. In our structure, pocket I interacts with the hydrophobic Ile instead of Pro at the 0 position) and pocket II accommodates Pro at the +3 position (Fig. 3A, 4D) from the gAnkB peptide. Prominently, ASAP1-SH3 contains two highly negatively charged pockets (pockets III & IV), with pocket III interacting with Lys at the +2 position and pocket IV accommodating Arg at the +5 position of the gAnkB peptide (Fig. 3A, 3D & 4D). Most uniquely, ASAP1-SH3 contains a large and flat negatively-charged surface that interacts electrostatically with Arg at the +6 position and likely also with Lys at the +7 position (Fig. 3A). These charge-charge interactions underlie the strong and specific interaction between gAnkB and ASAP1-SH3.

To validate our structural model, key residues in the gAnkB peptide were mutated to assess their effects on the binding to ASAP1-SH3. As summarized in Figure 3E, mutation of K1702 at the +3 position to Glu weakened the interaction by ∼120-fold; reverse the charge of R1705 at +6 position by Glu also weakened the interaction by ∼125-fold; a double mutation by replacing K1702 and R1705 with Glu completely abolished the interactions. These results support that the charge-charge interactions are critical for the binding of ASAP1 SH3 to the gAnkB peptide. Surprisingly, mutation of P1703 at the +4 position by a flexible Gly only decreased the binding affinity by ∼3-fold, indicating the sole Pro residue in the gAnkB peptide only plays a minor role for the ASAP1 SH3 binding. Taken together, the above structural and biochemical studies reveal that the ASAP1 SH3 domain recognizes gAnkB chiefly via extensive charge-charge interactions instead of the canonical “P-X-X-P“ motif-based bindings.

### Mechanisms underlie the high affinity and specificity of the ASAP1-SH3/gAnkB interaction

To elucidate the mechanism underlying the high affinity interaction between ASAP1-SH3 and gAnkB, we compared our structure with previously reported ASAP1-SH3/APC (PDB: 2rqu) and ASAP1-SH3/MICAL1 (PDB: 8hlo) structures. Similar to ASAP1-SH3/MICAL1 and ASAP1-SH3/APC structures, ASAP1-SH3 contains two additional highly negatively charged pockets (pockets III & IV) for incorporating the conserved lysine residue at +2 position and arginine residue at +5 position, in addition to the two SH3 domain signature pockets (pockets I & II) (Fig. 4A, 4B). Substantial differences in their binding mode also exist. Instead of a proline residue of the canonical “P-X-X-P“ motif in MICAL1 or APC (SAMP1 sequence), a highly conserved Ile at the 0 position in gAnkB is responsible for interacting with pocket 1 on ASAP1-SH3 (Fig. 3A, 4B & 4D). MICAL1 completely lacking the positively charged residues at the +6 and +7 positions to interact with a negatively charged flat surface unique to ASAP1-SH3 (Fig. 4B, 4D), explaining the weaker binding affinity of MICAL1 to ASAP1. APC has only one positively charged Lys at the +6 position and this Lys partially occupies the negatively charged flat surface of ASAP1 SH3 and the rest of the APC peptide turns away from the charged surface (Fig. 4B, 4D), again providing an explanation to the weaker binding of APC to ASAP1 ^45^. Interestingly, the residue at the +3 position of APC is a Ser instead of Pro (Fig. 4D) and this Ser residue can also occupy pocket II of ASAP1-SH3 (Fig. 4D), consistent with our result showing that the Pro at the +3 position in gAnkB peptide is not critical for the ASAP1 binding (Fig. 3E). In summary, the gAnkB peptide make full use of all binding sites (pockets I∼IV and the flat negatively-charged surface) on ASAP1-SH3 domain and thus binds to ASAP1 with a high affinity.

**Figure 4.**
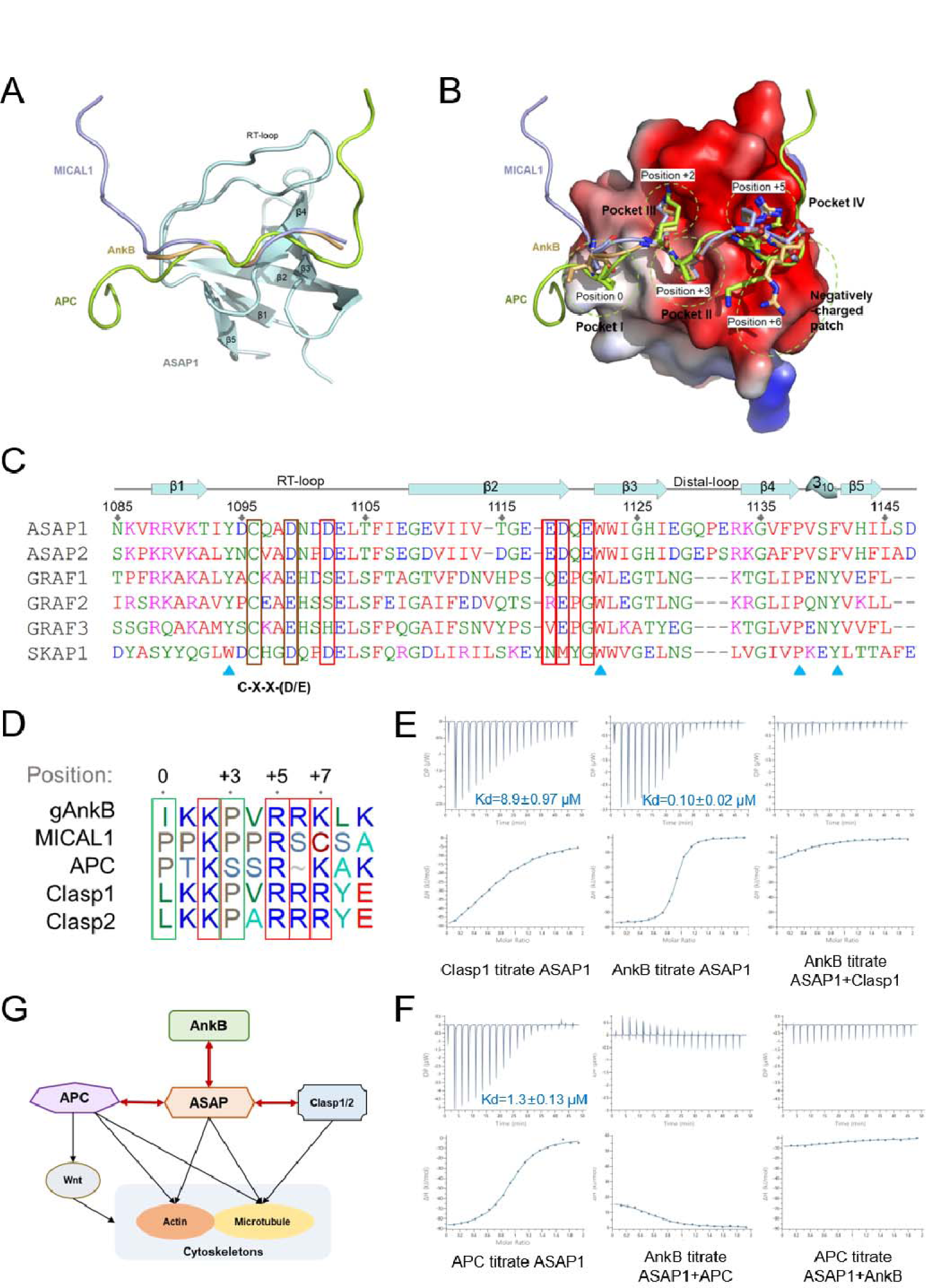
Mechanisms underlying the high affinity binding between ASAP1-SH3 and gAnkB. A: The ASAP1-SH3/gAnkB structure is aligned with the ASAP1-SH3/APC (PDB: 2rqu) and ASAP1-SH3/MICAL1 (PDB: 8hlo) structures. As SH3 domains from these structures are almost identical, for simplify, only the SH3 from the ASAP1-SH3/gAnkB structure is shown. B: Surface charge potential model combined with the stick model showing the bindings of the gAnkG, APC and MICAL1 peptides to ASAP1-SH3. The sidechains of the residues from the peptide ligands binding to pockets I∼IV are drawn in the stick model. C: Sequence alignment of six SH3 domains that contain the “C-X-X-D/E” motif in the RT-loop. Negatively charged residues in ASAP1-SH3 that are essential for binding to gAnkB are boxed in red. Key residues forming the pocket I& II are pointed by blue triangles (see Fig. 3C for their positions on the structure). D: Sequence alignment of the known ASAP1-SH3 binding ligand peptides including those from Clasp1/2 identified in the current study. APC shows the first SAMP repeating sequence region (SAMP1). Arbitrarily assigned positions of the motif are displayed on the top. E: ITC curves showing the bindings of CLASP1 or gAnkB to ASAP1-SH3, and the competition of gAnkB and Clasp1 in binding to ASAP1-SH3. F: ITC curves showing the binding of APC-SAMP2 to ASAP1-SH3 and the competition of gAnkB and APC-SAMP2 in binding to ASAP1-SH3. G: A hypothetic diagram showing the relationship of ASAP1 in linking gAnkB, APC and Clasp1/2 in regulating cytoskeletal structures in axons.

Similar to what was revealed by Jia et al. ^45^, a unique “C-X-X-(D/E)” sequence in the R/T- loop of ASAP1-SH3 creates a negatively charged binding pocket (pocket III) for the lysine residue at position +2 of the gAnkB peptide. This C-X-X-(D/E) sequence is present in ASAP1, ASAP2, GRAF1, GRAF2, GRAF3 and SKAP1, but not in the SH3 domains of other proteins. Comparing the sequences of the SH3 domains in these proteins revealed a series of residues in ASAP1-SH3 essential for gAnkB binding but partially or totally missing in GRAF1, GRAF2, GRAF3 or SKAP1 (Fig. 4C). The aspartic acid D1102 in the R/T-loop is in pocket IV and critical for binding to Arg (+5 position) (Fig. 3D). This residue is missing in the GRAF family proteins (Fig. 4C).

E1118 of ASAP1 at the end of the β2 strand also forms part of the pocket IV (Fig. 3D). This residue is missing in SKAP1 and the GRAF family SH3 domains (Fig. 4C). Finally, D1119 and E1121 of ASAP1, located in the loop between β2 and β3, together form the negatively charged flat surface patch interacting with the positively charged residue at the +6 and +7 positions of gAnkB.

These two negatively charged residues are partially or totally missing in SKAP1 and the GRAF family SH3 domains (Fig. 4C). Taken together, the above structure-based amino acid sequence analysis uncovers the molecular basis underlying the highly specific target recognition mechanism by the ASAP1/2 SH3 domains.

### Discovery of Clasp1/2 as ASAP1-SH3 binding proteins

To uncover potential ASAP1 binding proteins containing a motif similar to gAnkB, a search was conducted in the SWISSPROT protein sequence database for the [IVLAP]-X-K-P-X-R-[RK]- [RK] motif. Although most of them are not conserved during evolution, 197 such motif-containing sequences including AnkB, Clasp1, and Clasp2 were identified. The “ALKKPVRRRYE” sequence motif are highly conserved in Clasp1 or Clasp2 (Fig. S1B) and the motif is predicted to reside in the unfolded region of Clasp1 or Clasp2 by AlphaFold2. Clasp1 and Clasp2 are microtubule plus-end tracking proteins that regulate microtubule dynamics ^46–48^. ITC experiments using purified proteins confirmed that Clasp1 can bind to ASAP1-SH3 with Kd ∼8.9 μM (Fig. 4E). As expected, Clasp1 and gAnkB competed with each other in binding to ASAP-SH3 as the two proteins share a highly similar ASAP1-SH3 binding motif (Fig. 4A, 4D & 4E).

We also dissected the interaction between APC and ASAP1, Three sequence fragments (SAMP repeats) in APC were reported to bind ASAP1^49^, but no quantitative affinity data were available. We determined their affinity towards ASAP1 and found that the second SAMP repeat has the strongest affinity (Fig. S1C). This SAMP repeat sequence can also be nicely aligned with the ASAP1-SH3 binding motifs from gAnkB and Clasp1/2 (Fig. S1). As expected, APC also competed with gAnkB in binding to ASAP1-SH3 (Fig. 4F). Similar to Clasp1/Clasp2 and gAnkB, APC is also a cytoskeletal regulator, capable of directly regulating actin nucleation or microtubule remodeling ^50–56^. Both APC and Clasp proteins are involved in axon development. Thus, gAnkB, APC and Clasp1/2 may all participate in the regulation of axonal cytoskeletal structures in developing and/or mature neurons, possibly through binding to ASAP1 (Fig. 4G), a hypothesis that needs to be tested by future studies.

## Discussion

gAnkB is generated by acquiring an excessively long (∼2100 aa) and disordered sequence (neuron specific domain) during the evolution. A paralog of gAnkB, gAnkG also independently evolve to acquire an extremely long (∼2500 aa) intrinsically disordered insertion. Both giant ankyrins are specifically localized in axons, but they display distinct sub-axonal compartment localizations. To the best of our knowledge, all currently known binding partners for gAnkG also interact with gAnkB ^8,57–60^. It is puzzling how these two giant ankyrins are located in distinct axonal sub-compartments. Our current study uncovers a strong and direct interaction between the SH3 domain of ASAP1 and a short peptide from gAnkB, thus supporting a possible role of ASAP1 in axons ^61^. Importantly, this ASAP1-SH3 binding motif is absent in gAnkG, indicating that the two axon-localized giant ankyrins display distinct binding properties to ASAP1. Thus, our study begins to elucidate why the two giant ankyrins can play distinct functions in different axonal sub-compartments. It should be noted that we only covered 250 amino acid residues of the long insertion of gAnkB in our affinity purification experiment. It is possible that many other proteins may specifically bind to the long insertion of gAnkB or gAnkG, thus further defining their distinct functions in neuronal axons. Identification of such specific giant ankyrin binding proteins represents future fertile research grounds.

Both gAnkB and ASAP1 are cytoskeleton regulators. Cytoskeletons of axons are highly dynamic during development, but become extremely stable in mature neurons. For example, the formation of axon collateral branches is initiated by bundled actin filament-based axonal protrusions (filopodia) followed by dynamic microtubules invasion. Further stabilization of the invaded microtubules allow maturation and further extensions of these axonal branches ^62^. The whole axon branching process involves tight coupling of actin and microtubule dynamic regulation ^63–66^. gAnkB is a potent inhibitor of stochastic collateral axon branching by its ability to regulate microtubule dynamics and coupling microtubules with axonal plasma membranes, defects in which contributes to the formation of aberrant neuronal connections manifested in the autism models of rodents ^2,8^. Similarly, ASAP1 can directly bind to cytoskeletons and lipid membranes ^18,30,67^, though the roles of such interactions in axons have not been investigated. Moreover, ASAP1 may function through regulating the activity of ARF GTPases, which has been implicated in the regulation of membrane trafficking ^68–70^ and cytoskeletal remodeling ^71,72^. ASAP1’s reported binding partner APC as well as Clasp1/2 identified in this study all exist in axons. All these proteins are known to regulate actin and/or microtubule structures. Thus, ASAP1 may function as a hub in organizing these axonal cytoskeletal regulatory proteins including gAnkB, APC, and Clasp1/2.

## Materials and methods

### Constructs, protein expression and purification

The coding sequences of the ASAP1 constructs were PCR amplified from a mouse brain cDNA library. The coding sequences of AnkB constructs were PCR amplified from the full-length human 440-kD AnkB (a gift from Dr. Vann Bennett, Duke University). Clasp1, Clasp2, and APC coding sequences were PCR amplified from mouse brain or human HEK293T cDNA libraries. All of the constructs used for protein expression were cloned onto a home-modified pET32a vector with or without 3×StrepII tag. All truncation mutations or point mutations were produced with the same strategy as described in our previous study ^73^. Protein expression and purification protocols are the same as previously described ^73,74^. Recombinant proteins were expressed in BL21 (DE3) codon-plus Escherichia coli cells with induction by 0.1 mM IPTG at 16 LC. The N-terminal Trx- His_6_-tagged proteins were purified using Ni^2+^-NTA agarose affinity column followed by size- exclusion chromatography (Superdex 200 column from Cytiva) in the final buffer containing 50 mM Tris-HCl, pH 7.8, 1 mM DTT, and 1 mM EDTA, 100 mM NaCl.

### Affinity purification and Mass spectrum identification

The mouse brain specimens were obtained in accordance with the ethical review of laboratory animal welfare at Shenzhen Bay Laboratory. The brain tissue was immediately homogenized in a buffer solution consisting of 50 mM HEPES (pH 7.2), 600 mM NaCl, 15% glycerol, 0.1% Triton-X100, 20 mM CHAPS, 1 mM EDTA, 1 mM EGTA, 1 mM DTT, and a protease inhibitor cocktail (Cat. No. P8340, Sigma-Aldrich). The homogenates were then subjected to centrifugation at 90,000g for 40 minutes using an Ultracentrifuge (Optima XPN 100,

Beckman Coulter). The resulting supernatant was dialyzed in a buffer solution containing 20 mM HEPES (pH 7.2), 200 mM NaCl, 5% glycerol, 1 mM EDTA, 1 mM EGTA, and 1 mM DTT, followed by another round of ultracentrifugation at 100,000g for 40 minutes. The protein concentration in the supernatant was determined using the Bradford assay and the sample was prepared for the affinity purification.

For the affinity purification, the bait protein (gAnkB 1476-1727) was purified as a Trx-His_6_- 3×StrepII tag fusion protein, and conjugated to the StrepTactin Sepharose beads (Cytiva). Brain lysate (1 ml for each sample) was firstly precleared by StrepTactin beads and then mixed with the bait protein conjugated beads (20 μL dry volume for 1ml lysate), and incubated for 15 minutes, then centrifuged at 1000g for 1 min. The pellets containing the beads was washed three times with the dialysis buffer and then applied to SDS-PAGE. The SDS-PAGE gel was stained by Coomassie brilliant blue G-250 and protein bands were dissected for protein identification by Mass spectrometry by OmicSolution LTD., Shanghai, China.

### Pull-down and Western blot experiments

HEK293T cells transfected with Flag-tagged full-length ASAP1 plasmids were harvested 24 hours post-transfection. Cells were lysed in a lysis buffer containing 50 mM Tris pH 7.5, 100 mM NaCl, 1 mM DTT, 1% Triton X-100, and protease inhibitors (Sigma-Aldrich Cat. NO. P8340). The supernatant was obtained by centrifugation at 15,000g for 10 minutes at 4^◦^C, and then diluted four-fold with PBS. Purified gAnkB 1476-1727 protein (fused with Trx-His6-3×StrepII tag) was added to the diluted supernatant and incubated for 30 minutes. Afterwards, it was mixed with 20μL of StrepTactin beads and incubated for 10 minutes at 4^◦^C, followed by three washes with PBS buffer. Finally, the beads with bound proteins were mixed with loading buffer and boiled at 100 ^◦^C for 5 minutes, and detected by Western blotting using a mouse monoclonal anti-Flag antibody (Sigma-Aldrich, Cat. NO. F3165).

### Isothermal titration calorimetry (ITC) assay

Isothermal titration calorimetry (ITC) assays were carried out with the same protocol as described earlier using an ITC200 or a PEAQ-ITC MicroCal calorimeter (Malvern Panalytical) ^73^.

Briefly, in each titration a purified protein was loaded to the syringe (at 300 μM) and injected into the sample cell containing a binding partner (30 μM). The reaction temperature was set to 25 LC, each experiment contains 19 titration points and the first titration point was discarded for data analysis. Data were analyzed and fitted using the programs provided by the manufacturer.

### Crystallography

All crystals were obtained by sitting drop vapor diffusion methods at 16 LC. Crystals of gAnkB 1699-1710 in complex with ASAP1-SH3 were grown in solution containing 0.1 M MES monohydrate, pH 6.0, 40% v/v (+/-)-2-Methyl-2,4-pentanediol or 0.1 M MES monohydrate, pH 6.0, 20% w/v Polyethylene glycol 6,000. Crystals were soaked in crystallization solution containing additional 25% glycerol for cryoprotection. X-ray diffraction data were collected at BL02U1 beamlines at Shanghai Synchrotron Radiation Facility (SSRF) at 100 K. Data were processed and scaled using DIALS ^75^. Structures were solved by molecular replacement using PHASER ^76^ with the structure of ASAP1 SH3 in complex with the APC SAMP1 motif (PDB: 2rqu) as the searching model. Peptides were manually built according to the Fo-Fc difference maps in COOT ^77^. Further manual model adjustment and refinement were completed iteratively using COOT ^77^ and Refmac5 ^78^. The final models were validated by MolProbity ^79^ and statistics are summarized in Table 1. All structure figures were prepared by PyMOL (http://www.pymol.org).

## Acknowledgements

We thank the BL19U1 beamline at National Facility for Protein Science Shanghai (NFPS) for X-ray beam time. This work was supported by a grant from the National Nature Science Foundation of China (32000874) and grants from the Shenzhen Bay laboratory (S201101002, 21320011 and QH32001).

---

**Supplementary Figure 1:**
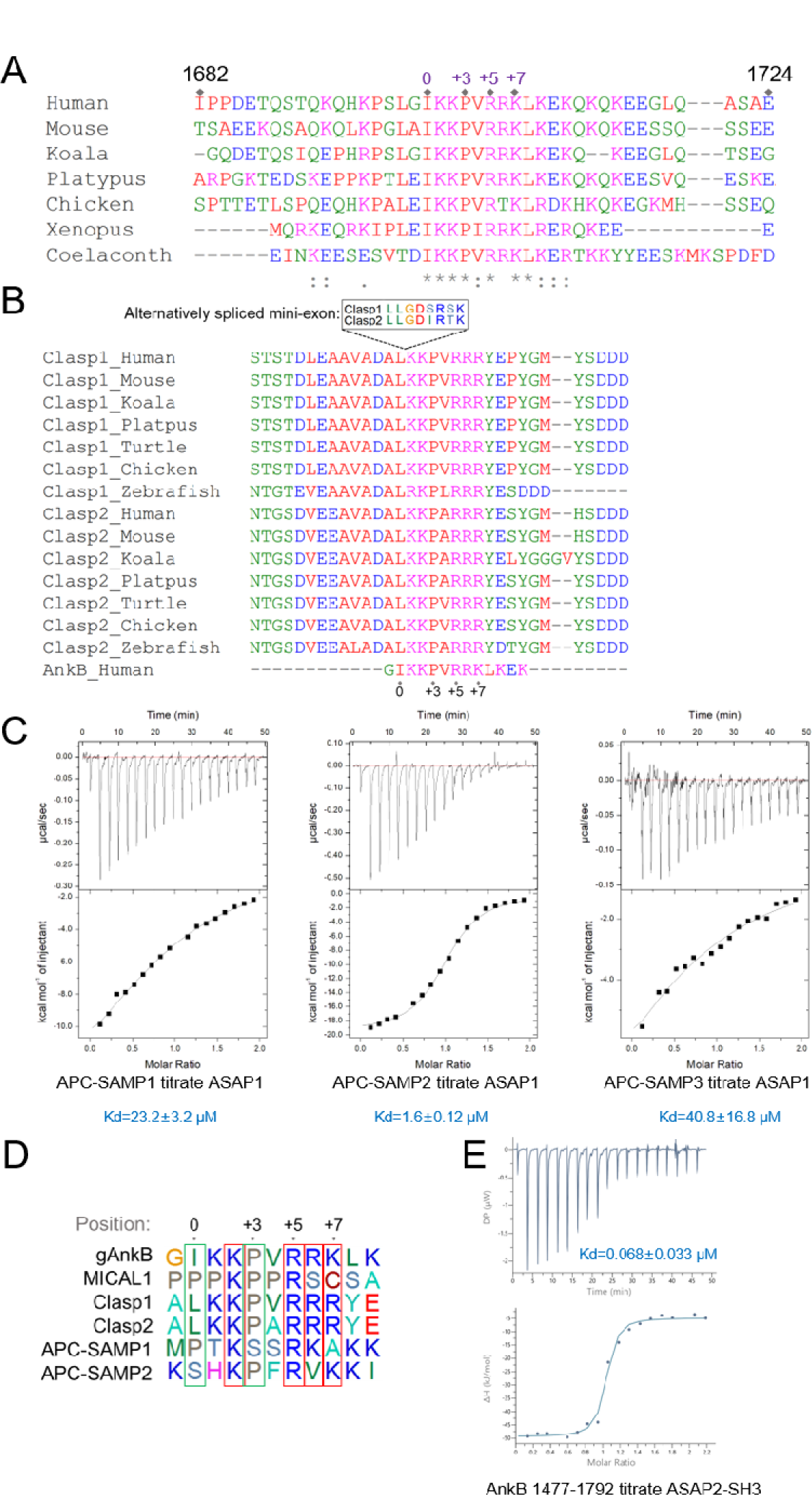
Clasp1/2 as newly identified ASAP1 SH3 binding proteins A: Alignment of the ASAP1-SH3 binding sequences from gAnkB across species. B: Alignment of the ASAP1-SH3 binding sequences from Clasp1, Clasp2 across species. The ASAP1-SH3 binding sequences from gAnkB is also included in the alignment. It is noted that Clasp1/2 contains an alternatively spliced mini-axon at the junction between Pro(0) and Lys(+1) in the ASAP1-SH3 binding motif. The Clasp1/2 isoform containing this mini-exon encoding sequence is likely to weaken or abolish the binding to ASAP1-SH3. C: ITC curves showing the interactions between three different SAMP repeat containing sequences from APC with ASAP1-SH3. APC-SAMP1, aa: 1546-1597; APC-SAMP2, aa: 1705-1754; APC-SAMP3, aa: 2009-2074. D: Sequence alignment of ASAP1-SH3 binding ligand peptides including APC-SAMP1 and APC- SAMP2. Arbitrarily assigned positions of the motif are displayed on the top. E: ITC-based assay showing that ASAP2-SH3 also binds to gAnkB with a very high affinity. The ITC curve shows the binding of gAnkB aa. 1477-1792 to mouse ASAP2-SH3 (aa. 896-958) recombinant protein.

